# Probability-based sequence comparison finds pre-eutherian nuclear mitochondrial DNA segments in mammalian genomes

**DOI:** 10.1101/2025.03.14.643190

**Authors:** Muyao Huang, Martin C. Frith

## Abstract

The insertion of mitochondrial genome-derived DNA sequences into the nuclear genome is a frequent event in organismal evolution, resulting in nuclear-mitochondrial DNA segments (NUMTs), which serve as a significant driving force for genome evolution. Once incorporated into the nuclear genome, some NUMTs can be conserved for extended periods and may potentially acquire novel cellular roles. However, current mainstream methods for detecting NUMTs are inefficient at identifying ancient and highly degraded NUMTs, leading to their prevalence and impact being underestimated. These ancient NUMTs likely play a far greater role in genetic functions than previously recognized, including contributing to the acquisition of functional exons. This study focuses on identifying ancient NUMTs in mammalian genomes using enhanced high-sensitivity sequence comparison methods. A sensitive and accurate NUMT-searching pipeline was established, predicting 1,013 NUMTs in the human reference genome, 364 (36%) of which are newly detected compared to the UCSC reference human NUMTs database. Notably, 90 pre-eutherian human NUMTs were identified, representing significantly older NUMTs than previously reported, with origins dating back at least 100 million years. The most ancient mammalian NUMTs could even date back over 160 million years, inserted into the nuclear genome of the common ancestor of therian mammals. This study provides a comprehensive exploration of the quantity and evolutionary history of mammalian NUMTs, paving the way for future research on endosymbiotic impact on the evolution of nuclear genomes.

## 1 Introduction

The endosymbiotic theory posits that mitochondria in modern animals and plants originated from the invasion of ancestral eukaryotic cells by Alphaproteobacteria (Margulis, 1970). Subsequent transfers of genetic material from the organelle to the nucleus ultimately shaped the mitochondria observed today, with many of their original genes integrated into the nuclear genome during the early stages of organelle evolution (Kleine et al., 2009; Perna and Kocher, 1996). Although the relocation of functional mitochondrial genes to the nuclear genome has become rare in recent evolutionary times (Boore, 1999; Kleine et al., 2009), the transfer of mitochondrial DNA (mtDNA) fragments to the nucleus remains an ongoing and frequent process across most eukaryotic lineages, giving rise to nuclear mitochondrial DNA segments (NUMTs). The precise mechanism by which NUMTs integrate into the nuclear genome is not yet fully understood. However, it is widely accepted that mtDNA fragments may be passively inserted at sites of nuclear double-strand breaks through nonhomologous end-joining, among several proposed mechanisms (Blanchard and Schmidt, 1996). Recent advances in sequencing technologies have enabled genome-wide surveys of NUMTs across diverse eukaryotic species, including both animals and plants (Calabrese et al., 2017; Liang et al., 2018; Zhang et al., 2020; Triant and Pearson, 2022). The number and size distribution of NUMTs vary considerably among species. While some animal genomes harbor only a few or no detectable NUMTs, others contain thousands, ranging in size from just a few base pairs to several hundred kilobases (Hazkani-Covo et al., 2010; Hazkani-Covo, 2022). NUMTs are often regarded as “dead-on-arrival” pseudogenes. Once inserted into the nuclear genome, most NUMTs begin to degrade, losing their original functions and undergoing various post-integration fates, including elimination, mutation, proliferation, rearrangement, and fragmentation (Zhang et al., 2020). These post-integration processes obscure the boundaries and timing of NUMT insertion events, rendering their identification particularly challenging. Such complexities are especially pronounced for ancient NUMTs, whose substantial sequence divergence from the mitochondrial genome necessitates more sensitive and accurate detection approaches.

As a significant driving force influencing the evolution of the nuclear genome, NUMTs were considered to be non-functional and harmless for a long time, mainly contributing to the birth of non-coding sequences and pseudogenes. However, as investigations into NUMTs become increasingly comprehensive, more potential impacts of NUMT insertions are being uncovered. For instance, NUMT insertions have been reported to disrupt normal gene expression and are thought to contribute to various human diseases, including rare genetic disorders (Borensztajn et al., 2002; Turner et al., 2003) and cancers (Ricchetti et al., 2004; Puertas and Gonźalez-Śanchez, 2020; Wei et al., 2022), potentially through interference with tumor suppressor genes or the activation of oncogenes. In addition to their recent insertion into genes that disrupts normal expression, NUMTs have also been recruited as novel exons, contributing new coding sequences to preexisting nuclear genes (Noutsos et al., 2007). It has been proposed that NUMTs may have contributed to more ancient functional exon acquisitions, which were previously difficult to detect due to the limitations of conventional sequence comparisons. NUMTs have also been utilized as phylogenetic markers, owing to their slower nuclear mutation rates and homoplasy-free insertions (Liang et al., 2018). However, their high similarity to mitochondrial DNA may result in misidentification, thereby complicating analyses such as heteroplasmy detection, ancient DNA studies, and population genetics (van der Kuyl et al., 1995; Zhang and Hewitt, 1996; Albayrak et al., 2016). Therefore, accurate and comprehensive identification of NUMTs is essential for understanding their evolutionary roles, minimizing their confounding effects, and ensuring reliable mitochondrial DNA analyses.

Current NUMT identification methods are generally categorized into wet-lab-based and computational approaches (Xue et al., 2023). Computational methods typically target either ancestral NUMTs present in the reference genome or rare, polymorphic NUMTs observed in individuals. While advances in next-generation sequencing have facilitated the detection of recent insertions, the annotation of ancestral NUMTs in the reference genome has lagged behind. Databases of common human NUMTs remain outdated, with annotations still based on genome version hg19. Nevertheless, accurate annotation of ancestral NUMTs remains essential for both excluding them during polymorphic NUMT detection and advancing evolutionary studies, which often fail to detect ancient insertions due to prior methodological limitations. In previous studies, NUMTs have typically been identified by aligning the mitochondrial genome to the nuclear genome using local sequence alignment tools (Simone et al., 2011; Tsuji et al., 2012; Uvizl et al., 2024), with BLASTN being one of the most widely used tools. However, the results are sensitive to search parameters and thresholds. Moreover, BLASTN relies on fixed scoring schemes, which may reduce detection sensitivity. While DNA-to-protein alignments utilize informative protein scoring matrices, DNA-to-DNA alignments depend on a simplistic 4×4 scoring matrix, potentially contributing to BLASTN’s suboptimal performance in identifying ancient NUMTs. Recent work has highlighted the advantages of DNA-to-protein comparisons in improving the sensitivity of similarity searches. Protein sequences, due to their slower evolutionary rate and better scoring models, often enable more accurate homology inference over long evolutionary distances (Pearson, 2019). However, this approach fails to detect alignments involving non-coding mitochondrial sequences. A combined approach utilizing both DNA-to-protein and DNA-to-DNA comparisons has been proposed to enhance both sensitivity and accuracy in NUMT detection (Triant and Pearson, 2022); however, additional refinements remain necessary, particularly for the detection of ancient insertions, which often show reduced sequence similarity due to long-term mutational changes and other evolutionary factors.

Here, an optimized and accurate NUMT detection pipeline was developed based on LAST and integrates both DNA-to-protein and DNA-to-DNA matching strategies. A key advantage of the pipeline lies in its ability to learn scoring matrices and filtering thresholds tailored to the input sequences. Scoring parameters are optimized via maximum-likelihood estimation (Hamada et al., 2017), thereby enabling alignment to reflect sequence-specific patterns and evolutionary constraints, which enhances the accuracy of ancient NUMT detection. Moreover, the DNA-to-protein module allows frameshifts within matches and employs a 64×21 substitution matrix, rather than the standard 20×20 matrix, to improve sensitivity (Yao and Frith, 2022). Conventional DNA-to-protein matching approaches have relied on 20×20 amino acid substitution matrices, such as BLOSUM, which are derived from substitution rates in extant proteins and may be suboptimal for detecting protein “fossils” such as NUMTs (Frith, 2022). This enhancement facilitates the identification of subtle similarities, thus enhancing sensitivity in detecting NUMTs that have undergone significant sequence divergence. The pipeline is well-suited for detecting highly degenerated NUMTs. Using the optimized pipeline, NUMTs were annotated in the human reference genome hg38, identified in 15 additional mammalian reference genomes, and traced across mammalian clades, providing estimates of their insertion times.

## 2 Materials and Methods

### 2.1 Genome Data

16 annotated reference genomes, including mitochondrial and nuclear sequences from various mammalian orders, were downloaded from NCBI. The species analyzed represent 9 orders: Artiodactyla (2 species), Carnivora (3 species), Lagomorpha (1 species), Marsupial (2 species), Monotremata (2 species), Pilosa (1 species), Primates (3 species), Proboscidea (1 species), and Rodentia (1 species). Each genome was assembled to chromosome-level quality in its latest version (May 2024). Unplaced scaffolds were excluded from this study to ensure accurate NUMT identification. 13 mitochondrial proteins for each species were downloaded separately from NCBI. Details of the genome data, including species and assembly accessions, are provided in Supplementary Table S1.

### 2.2 NUMTs detection pipeline

NUMTs were detected using a novel detection pipeline based on LAST version 1639 (Kielbasa et al., 2011). The pipeline consists of four major components: initial alignment, reverse test, quality filtering, and integration.

#### 2.2.1 Initial alignments

In the first step, two types of sequence comparisons were performed: nuclear genome vs. mitochondrial genome (DNA-to-DNA), and nuclear genome vs. mitochondrial proteins (DNA-to-protein). For the DNA-to-DNA comparison, the nuclear genome was used as the query and the mitochondrial genome as the reference. LAST was applied to identify alignments across the entire nuclear genome by pre-estimating substitution and gap frequencies between the nuclear and mitochondrial sequences using `last-train` (Hamada et al., 2017), as illustrated by the following commands:

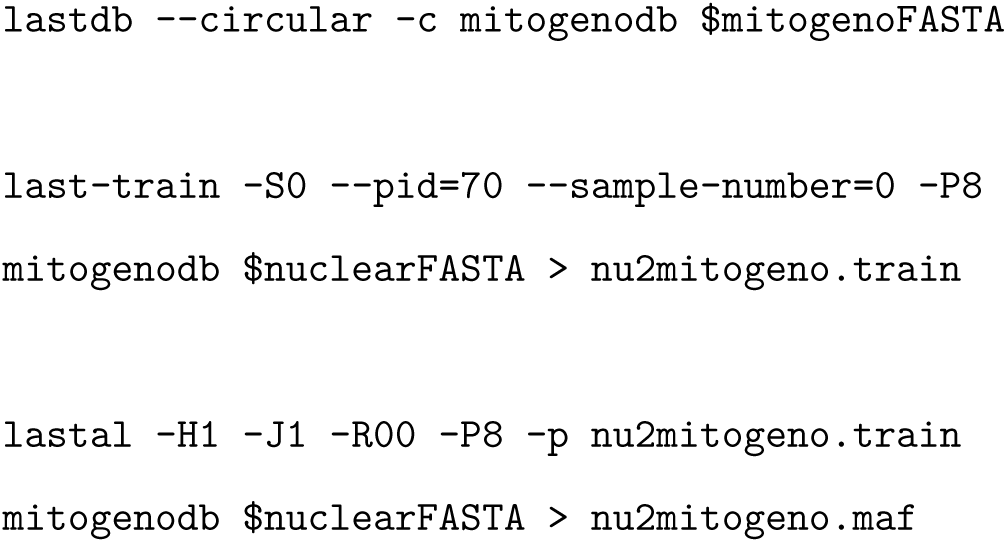

LAST commands in the DNA-to-protein comparison are shown below:

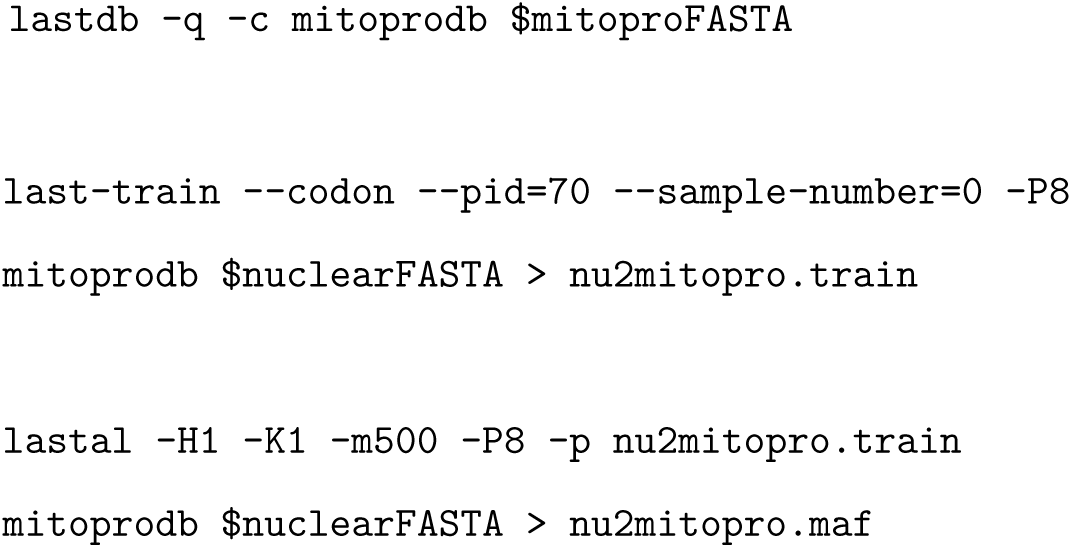

Several options were used in the LAST commands to improve sensitivity and ensure accuracy in detecting ancient (low-identity) NUMTs. In DNA-to-DNA comparisons, the `--circular` option enables the comparison to treat the mitochondrial genome as circular by appending a copy of each sequence to itself, thereby avoiding the loss of hits at sequence boundaries. The `-c` option, combined with `-R00`, soft-masks repetitive elements from mitochondrial sequences while retaining them in nuclear sequences during the initial matches. It helps prevent false homologous predictions while still allowing the detection of NUMTs with tandem duplicatiions in the nuclear genome. In `last-train`, the `--pid=70` option ignores alignments with *≥* 70% identity, enhancing the detection of ancient NUMTs at the expense of missing very short, high-identity NUMTs. This is consistent with the goal of identifying the oldest NUMTs within genomes. The `--sample-number=0` option ensures `last-train` uses the whole query sequence to train for the optimal scoring matrix. -`H1` option reports alignments that are expected to occur by chance at most once in all sequences. The `-J1` option uses a new approach to better detect subtly related sequences by summing the probabilities of all alternative alignments between them (Frith, 2024). `-P8` enables parallel processing with 8 threads to enhance computational speed. In DNA-to-protein comparison, the `-K1` option means that for each region of the nuclear genome, only the alignment with the highest score will be output. The `-m500` option makes the search slower but more sensitive.

#### 2.2.2 Reverse test

Following the initial alignment, preliminary hits from the two comparisons were respectively masked in the query sequence, which was then reversed (but not complemented). The reversed, masked query sequence was subsequently used to repeat both the DNA-to-DNA and DNA-to-protein comparisons, serving as a negative control. Masking was necessary because using an unmasked reversed sequence as a decoy would artificially elevate the background similarity, thereby overestimating the filtering threshold and potentially discarding true ancient NUMTs (Glidden-Handgis and Wheeler, 2024). Since genuine NUMTs are not expected to appear in the reverse test, any hits identified in this step are considered random background noise.

#### 2.2.3 Quality filtering

During quality filtering, the highest alignment score observed in the reverse test was adopted as a threshold, and all candidate alignments from the initial comparison with lower scores were discarded.

Additionally, as an important novel step, alignments overlapping with nuclear ribosomal RNA (rRNA) regions were excluded. Nuclear rRNA data for all species, except for humans, was collected using LAST. For humans, rRNA data were downloaded from the RepeatMasker track in the UCSC Table Browser, as the rRNA annotation datasets were available for the latest genome assembly used in this research. Since rRNA is known to be highly conserved in eukaryotes (Dalal and Lyons, 2023; Symonová, 2019), we accelerated the search for rRNA genes or pseudogenes in other genomes by comparing the nuclear genomes of the 14 species to the human rRNA gene RNA28SN5 using the following LAST commands:

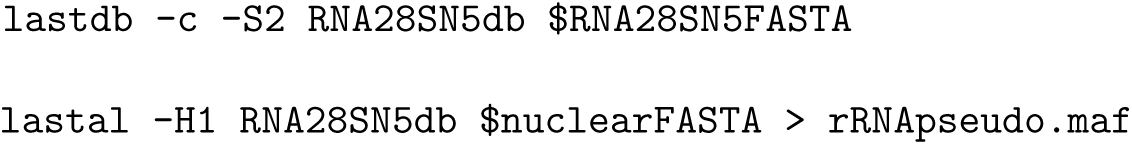

#### 2.2.4 Integration

The strand orientation in the DNA-to-protein comparison is not consistent with that in DNA-to-DNA comparison. Therefore, aligning the strand direction between the two sets of results is necessary for subsequent analysis. To achieve this, the strand of origin for each mitochondrial protein must be determined based on its position in the mitochondrial genome. This was performed by the following commands. `fix-protein-strand` checks the strand orientation of mitochondrial proteins in the mitochondrial genome and adjusts the strand information of alignments in the DNA-to-protein comparison accordingly. Detailed commands and implementation are available at: https://github.com/Koumokuyou/NUMTs/, which also hosts a UCSC Genome Browser track hub containing several processed genomic tracks and annotations generated by the NUMT detection pipeline described in this study.

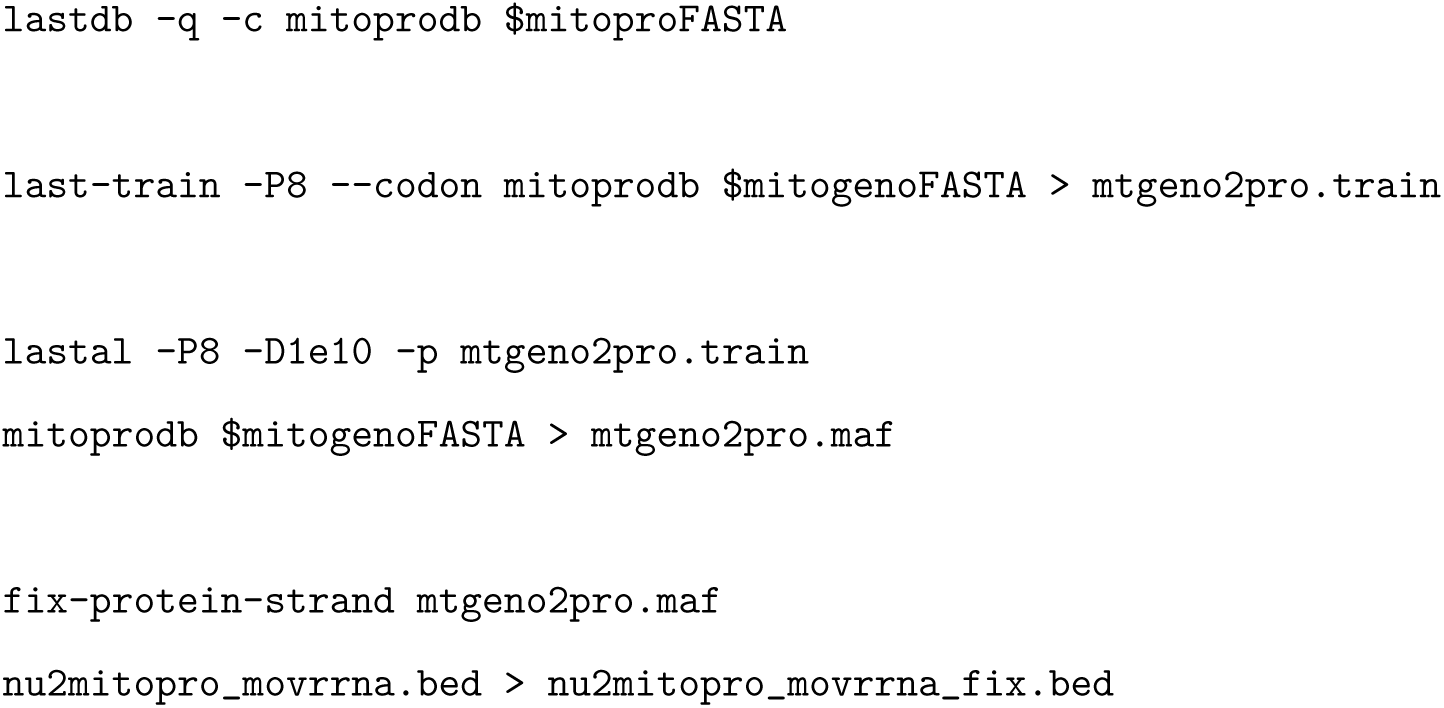

Finally, filtered alignments from the DNA-to-DNA and DNA-to-protein comparisons were merged using BEDTools v2.29.1 to generate the final NUMT dataset. Alignments from the two comparisons were merged into a single NUMT if they were on the same strand and either overlapped by at least 1 bp or were directly adjacent (i.e., bookended). Additionally, merged alignments shorter than 30 bp were excluded. This length threshold was empirically determined based on previous studies (Triant and Pearson, 2022; Uvizl et al., 2024) to ensure that retained sequences were sufficiently long to produce statistically significant alignments.

#### 2.2.5 Assembly of NUMT segments

To facilitate subsequent analyses for constructing the NUMT orthology network, adjacent NUMTs were grouped into larger genomic regions, hereafter referred to as blocks, using BEDTools. A block was defined as a genomic region consisting of one or more NUMT segments located within 2000 bp of each other in the nuclear genome, regardless of strand orientation (Uvizl et al., 2024). This strategy consolidated fragmented NUMT sequences into unified units, thereby simplifying the analysis of their evolutionary relationships.

### 2.3 Identification of ancient NUMTs in the mammalian genomes

#### 2.3.1 Pairwise alignments between species

120 pairwise genome alignments were conducted by LAST, ensuring that only orthologous segments were identified while excluding non-homologous insertions between any two species within the 16 species (Frith and Kawaguchi, 2015):

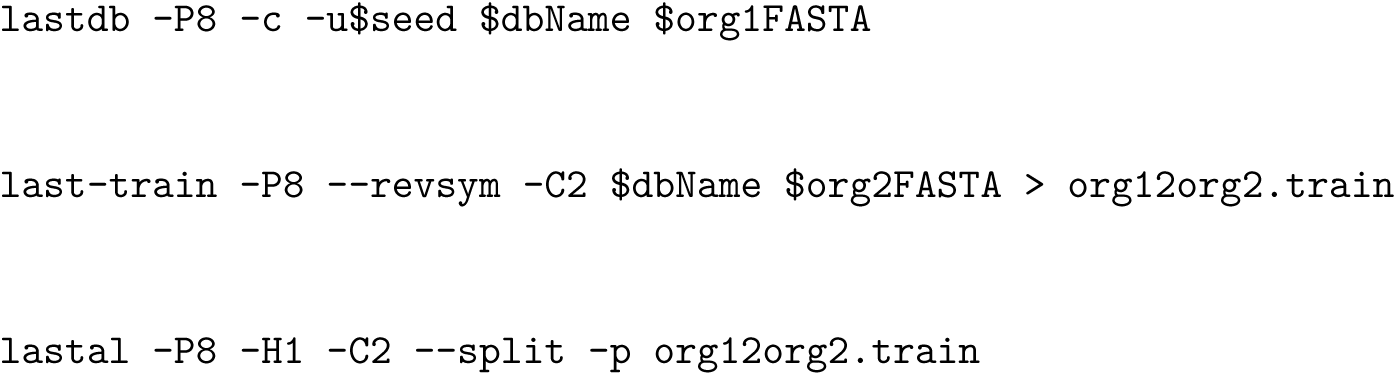
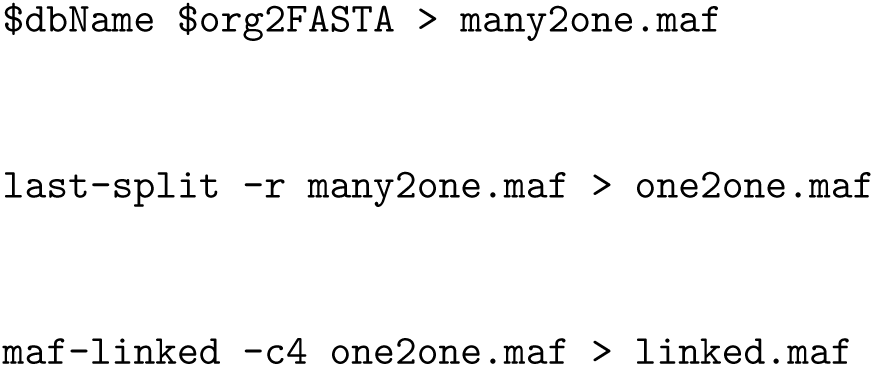

To obtain accurate orthologous alignments, the seeding scheme setting $`seed` depended on the relationship between two genomes. For two genomes that came from the same order, $`seed` was set to `RY4` (Frith et al., 2023); for two genomes that came from the same class but different orders, $`seed` was set to `YASS`. `maf-linked` was used to remove isolate alignments in genome-to-genome alignments with the aim of discarding alignments between non-homologous insertions of homologous segments, including NUMTs (Frith, 2022). Adjacent alignments, with a minimum of four, were ”linked” if they were within the distance of 10^6^ bp and separated by no more than five other alignments.

#### 2.3.2 Construction of ancient NUMT block repository

A NUMT identified in a given species was considered ancient if it overlapped with at least two pairwise genome-to-genome alignments between that species and other species. Based on this criterion, ancient NUMT blocks were identified across 16 species and compiled into species-specific repositories for subsequent analyses.

### 2.4 NUMT orthology relationships between species

The approach used to identify orthologous NUMTs was similar to that described by Hazkani-Covo et al. (Hazkani-Covo and Graur, 2007). Orthology relationships were inferred between species pairs based on the ancient NUMT block repositories constructed in the preceding step. For a given species pair (e.g., species A and species B), preliminary orthologous NUMTs were identified by assessing whether NUMT blocks from each species overlapped with pairwise genome alignments by at least 50% of their length. The subsequent classification of orthologous relationships was based on the nature of the alignment and the presence or absence of NUMTs in the aligned regions. Three types of orthology scenarios were defined:

1. **Type 1:** A NUMT block from species A overlapped with orthologous segments between species A and B and also with at least one NUMT block from species B within those segments. In this case, the NUMT blocks from both species were considered orthologous. The number of orthologous NUMTs between A and B was incremented by the number of species B blocks that overlapped with the species A block.
2. **Type 2:** A NUMT block from species A overlapped with orthologous segments between species A and B, but no NUMT block from species B was present in the aligned region. This absence suggested the possible presence of a highly degenerated NUMT in species B that was not detected through standard comparisons. To account for this possibility, the aligned region in species B corresponding to the species A NUMT block was added to the species B ancient NUMT repository as a putative NUMT. This pair was included in the count of orthologous NUMTs.
3. **Type 3:** The inverse of Type 2. A NUMT block from species B overlapped with orthologous segments, but no NUMT block was detected in the corresponding region of species A. In this case, the aligned region in species A was added to the species A NUMT repository as a putative ancient NUMT block. The NUMT block from species B, along with its inferred counterpart in species A, was counted as an orthologous pair.

## 3 Results and Discussion

### 3.1 Optimized NUMTs searching pipeline

For each species, NUMTs were identified through comparisons between the nuclear genome and the mitochondrial genome, the mitochondrial proteome, or both. LAST makes efforts to find alignments for every coordinate in the nuclear genome by pre-learning the substitution and gap rates between the mitochondrial genome and the nuclear genome (Hamada et al., 2017). A sample scoring matrix trained on alignments between the human nuclear and mitochondrial genomes is shown in Figure 2a, while the matrix trained on alignments between mitochondrial proteins and the nuclear genome is shown in Figure 2b. It is noteworthy that the human mitochondrial genome comprises the heavy (H)-strand and light (L)-strand and data from NCBI provides the L-strand sequence of the mitochondrial genome. Our comparisons and learned scoring matrix are based on the L-strand mitochondrial sequence.

**Figure 1:**
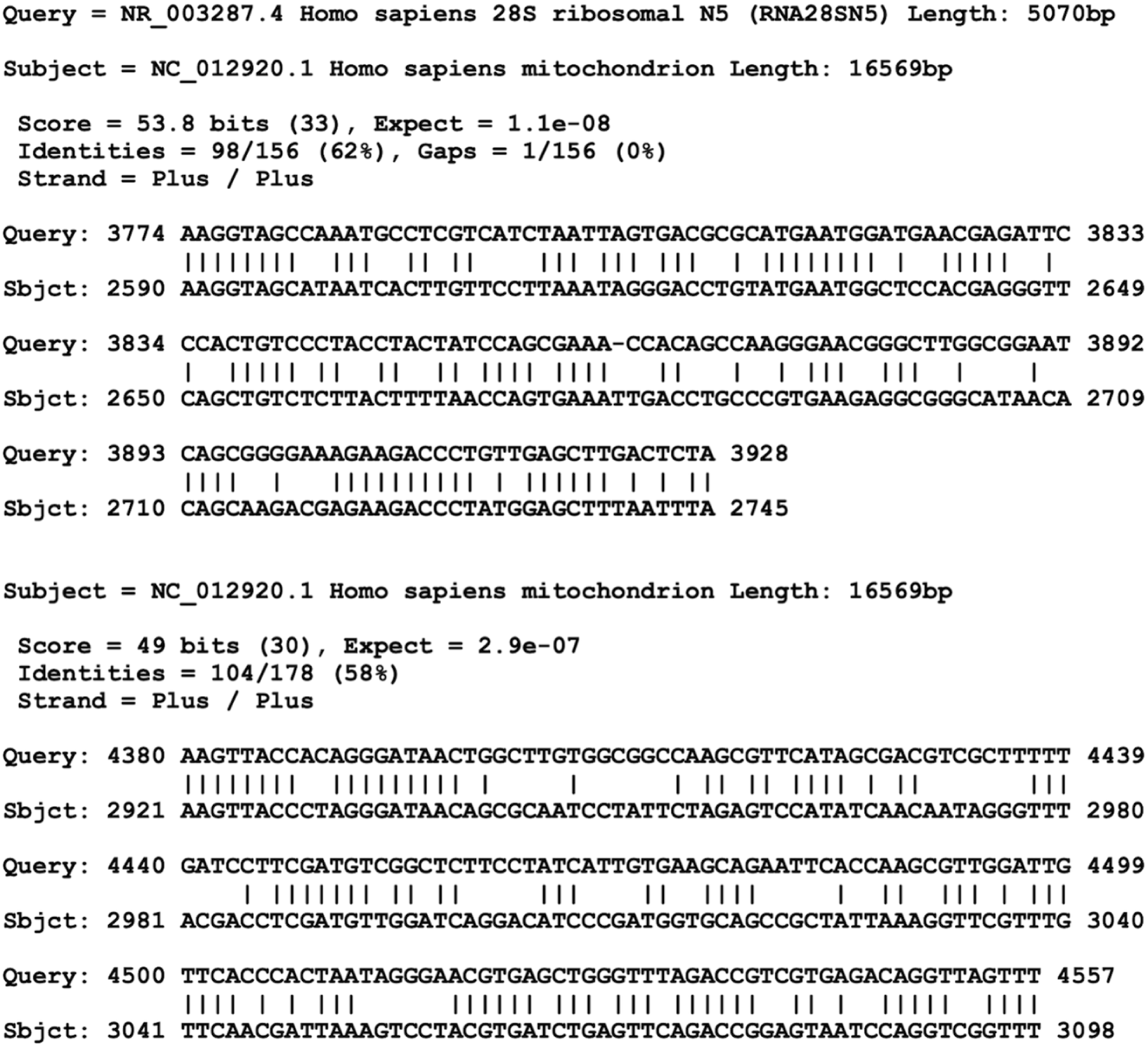
Alignments between the human mitochondrial genome and the nuclear-encoded ribosomal RNA gene RNA28SN5 identified using the LAST alignment tool. The query sequence is the human nuclear 28S rRNA gene (RNA28SN5), and the reference sequence is the human mitochondrial genome (NC 012920.1). Each alignment presents nucleotide-level matches between the two sequences, with vertical bars indicating exact matches. The aligned regions in the mitochondrial genome are both located within the annotated 16S rRNA gene (mitochondrial large rRNA subunit). Alignment scores, percent identity, gap information, and strand orientation are indicated for each block. This figure was made with `maf-convert` from the LAST package.

**Figure 2:**
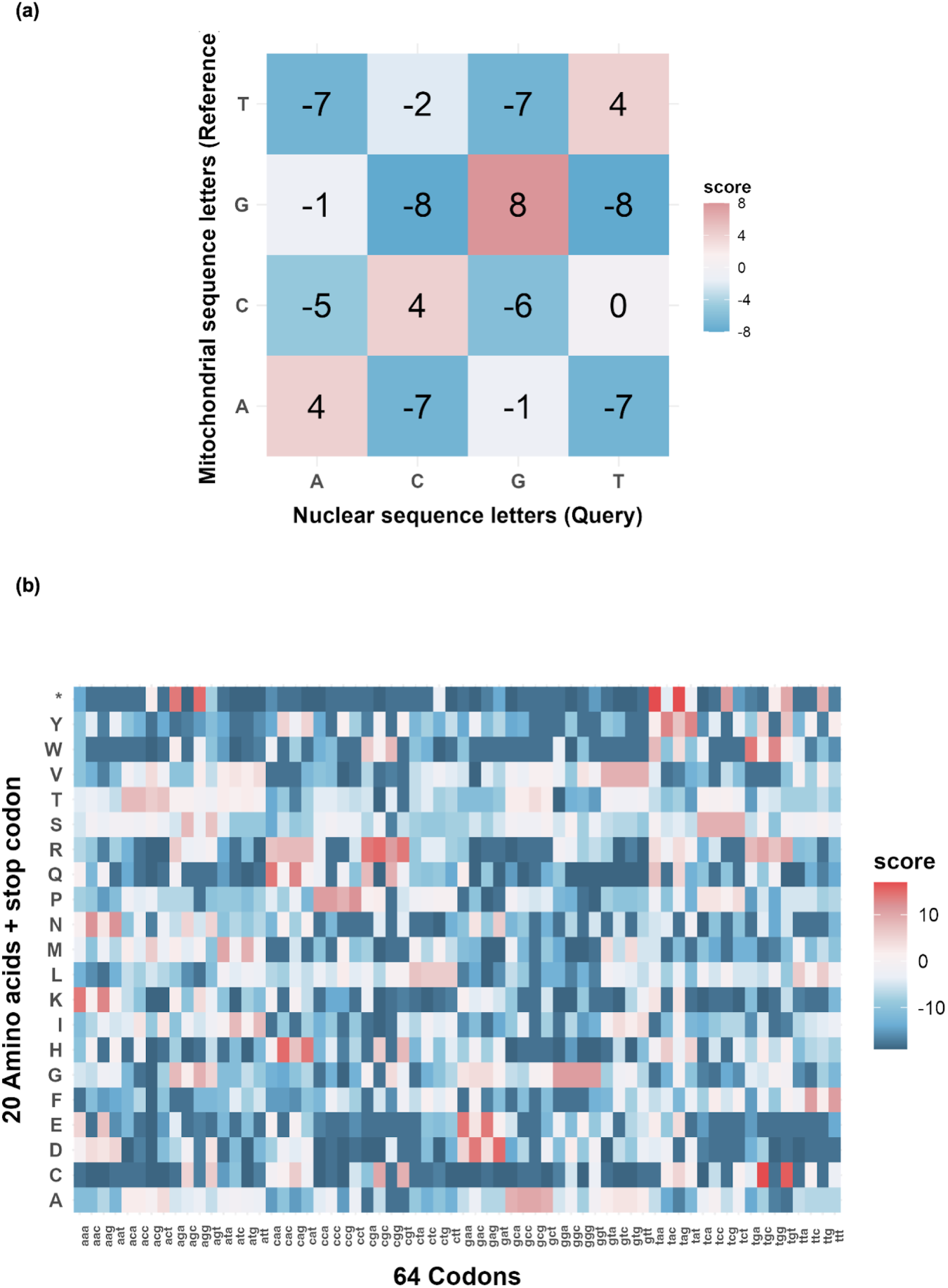
Substitution matrices learned from nuclear-mitochondrial sequence alignments in the human genome. (a) A 4×4 substitution matrix inferred from the human nuclear genome versus the mitochondrial genome. Positive values (e.g., G–G) indicate favored matches, while negative scores represent disfavored or unlikely substitutions. (b) A 64×21 substitution matrix (64 codons × 20 amino acids plus STOP), inferred from the human nuclear genome versus mitochondrial protein. In both heatmaps, the color gradient represents substitution scores, ranging from deep blue (strongly negative scores) to deep red (strongly positive scores), with white indicating neutral or near-zero substitution scores.

Using species-specific scoring matrices pre-trained with LAST, DNA-to-DNA comparisons typically recover the majority of NUMTs. However, extremely ancient mitochondrial protein ”fossils” that are highly diverged at the nucleotide level may be missed by DNA-only comparisons and can instead be effectively detected through DNA-to-protein alignments (Frith, 2022).

To estimate the false positive rate and reduce potential biases associated with arbitrarily defined thresholds, we applied a refined reverse test for each species. It has been reported that reverse alignments can yield artificially high alignment scores when palindromic structures are present, potentially leading to an overestimation of false matches when reversed sequences are used as decoys (Glidden-Handgis and Wheeler, 2024). This effect is particularly problematic when identifying ancient NUMTs, whose borderline E-values can be difficult to distinguish from background noise. To address this, we used a reversed version of the query sequence, with pre-identified NUMT regions masked, as a decoy for estimating background similarity.

Alignments overlapping with nuclear ribosomal RNA (rRNA) regions were excluded from the results. Due to LAST’s high sensitivity, spurious alignments often occur between the mitochondrial 16S rRNA gene and homologous nuclear rRNA sequences (Figure 1). These false alignments may inflate the estimated abundance of ancient NUMTs, as conserved nuclear rRNA regions can be mistakenly interpreted as mitochondrial-derived insertions.

To date, no standardized pipeline or universally accepted criteria exist for the detection of NUMT insertions. The most widely used local alignment tool is BLASTN, which requires users to manually select among several pre-defined alignment tasks and scoring matrices, and to set E-value thresholds themselves (Uvizl et al., 2024; Hazkani-Covo and Martin, 2017; Calabrese et al., 2012; Hazkani-Covo and Graur, 2007). To accommodate different detection objectives, researchers often adjust search parameters iteratively to optimize results. However, in the absence of a gold standard for NUMT identification, parameter inconsistencies across studies frequently lead to divergent NUMT sets, thereby complicating cross-study comparisons. Moreover, BLASTN, which typically uses extant mitochondrial nucleotide sequences as queries, is generally limited to the detection of relatively recent NUMTs (Mishmar et al., 2004). It performs poorly when attempting to recover ancient NUMTs, which tend to be highly fragmented and diverged due to extensive mutation over long evolutionary timescales. To address this limitation, previous studies have applied DNA-to-protein alignment tools such as TBLASTN and TFASTX, which improve sensitivity for detecting divergent NUMTs by utilizing protein-level conservation (Triant and Pearson, 2022; Antunes and Ramos, 2005). Despite their increased sensitivity, these methods still face difficulties in distinguishing true ancient NUMTs from spurious matches, particularly in low-complexity genomic regions. This challenge has contributed to conflicting results in earlier studies. For example, Antunes and Ramos (Antunes and Ramos, 2005) reported the detection of 5,621 ancient NUMTs in the Fugu genome (assembly v2.0) using TFASTX with the BLOSUM100 matrix. In contrast, Venkatesh et al. (Venkatesh et al., 2006) later argued that many of these were likely false positives, resulting from low-stringency search parameters and spurious alignments to repetitive nuclear sequences rather than genuine mitochondrial insertions. In response to these limitations, we developed a sequence-driven NUMT detection pipeline that employs species-specific scoring matrices and adaptive E-value thresholds, thereby reducing the subjectivity associated with manual parameter setting. By leveraging the accurate E-value estimation of LAST, our method effectively enhances the recovery of ancient NUMTs while minimizing false positives, offering a more robust and reproducible framework for cross-species NUMT discovery.

### 3.2 Newly detected human NUMTs

Using our optimized NUMT-searching pipeline, we identified a total of 1,013 NUMTs in the human reference genome (GRCh38/hg38). The number of NUMTs per chromosome ranged from approximately 20 to 100. To evaluate the accuracy of our predictions, we compared our dataset with the reference UCSC human NUMT database, which is based on the GRCh37/hg19 assembly (Simone et al., 2011). For comparison, the UCSC NUMT coordinates were converted to the hg38 assembly using the UCSC LiftOver tool, with 750 out of 766 entries successfully mapped. Among these 750 UCSC NUMTs, our pipeline recovered 737 (98.2%) with over 90% sequence overlap, 739 (98.5%) with over 50% overlap, and 742 (98.9%) with at least 1 bp overlap (Figure 3a). This high recovery rate demonstrates the robustness and reliability of our method in detecting previously annotated NUMTs. We further checked the eight UCSC-only NUMTs and found that five had indeed been detected by our pipeline and included in our UCSC Genome Browser submission, but were later filtered out during our strict quality control process.

**Figure 3:**
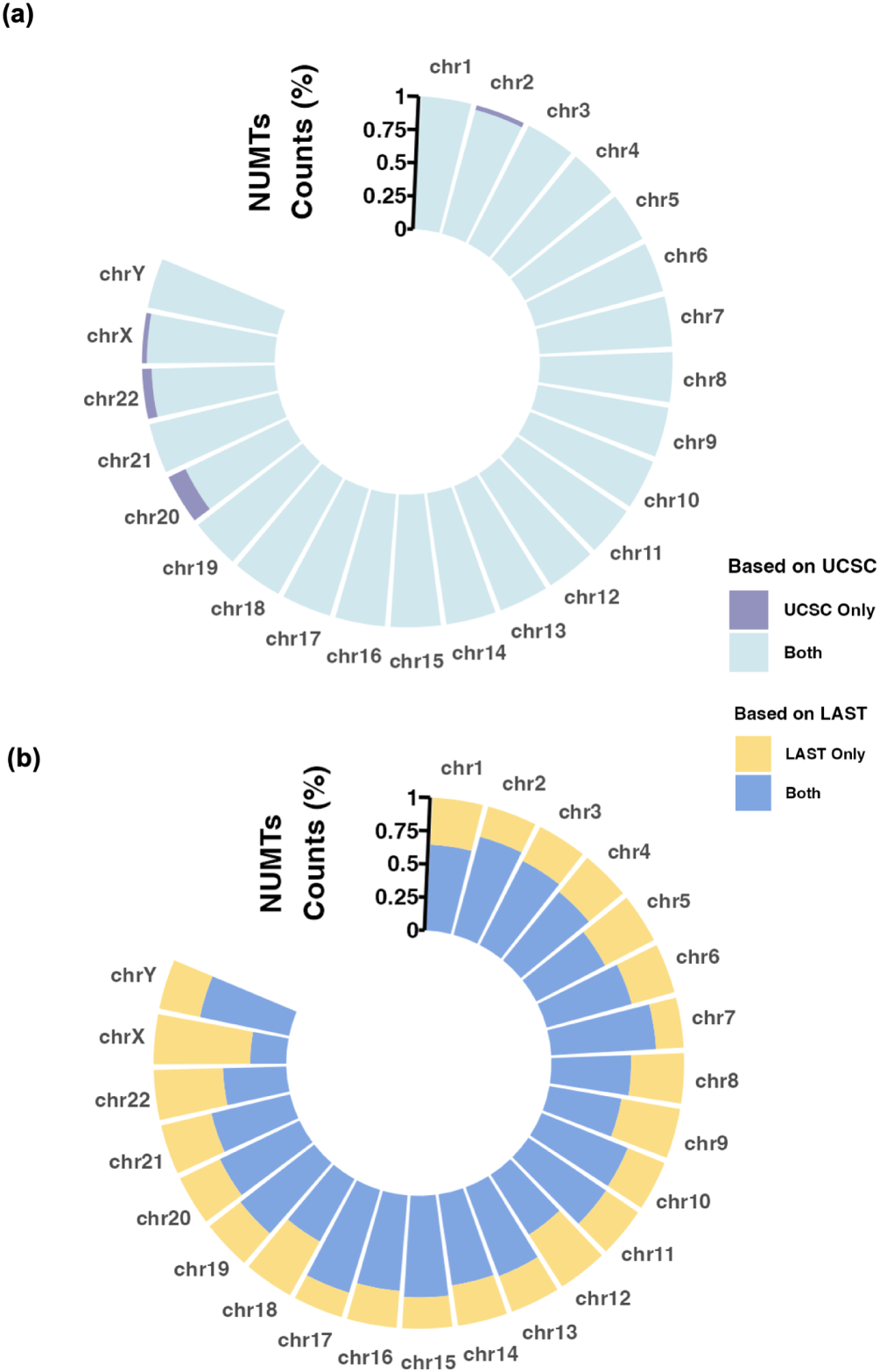
Comparison of NUMT distributions across human chromosomes between our pipeline and the UCSC 2011 reference database. (a) Proportional distribution of NUMTs per chromosome based on the UCSC NUMTs dataset (converted to hg38). Blue segments represent NUMTs that overlap (by at least one base) with entries in our dataset, while purple segments indicate UCSC-specific NUMTs not recovered by our method. (b) Proportional distribution of NUMTs per chromosome based on our NUMTs dataset. Blue segments indicate NUMTs overlapping with those in the UCSC dataset, while yellow segments represent NUMTs uniquely identified by our method. The comparison highlights both the shared and distinct genomic coverage between the two approaches.

In addition to recovering nearly all UCSC reported NUMTs, our pipeline identified 364 novel NUMTs not present in the UCSC database, accounting for 39% of the total predictions (Figure 3b). These newly discovered NUMTs were supported by stringent global E-value thresholds and reverse-alignment filtering, suggesting that they are true insertions rather than random matches. Their absence from the UCSC dataset highlights the enhanced sensitivity of our approach, particularly in detecting ancient or highly degenerated NUMTs that previous methods may have missed.

Among the remaining 649 NUMTs overlapping with UCSC-reported entries, we examined the type of comparison through which each was detected. We found that 218 (34%) were identified exclusively through DNA-to-DNA comparison, while 431 (66%) were detected in both DNA:DNA and protein:DNA comparisons. Notably, the UCSC dataset contained no NUMT detected exclusively through DNA-to-protein comparison. In contrast, our dataset included 42 such cases (4% of the total), underscoring the strength of our approach in capturing ancient mitochondrial protein ”fossils” that are often overlooked by nucleotide-based methods alone.

### 3.3 Comprehensive annotation of human NUMTs in hg38

Our hg38 NUMT dataset, formatted in BED, has been uploaded to the UCSC Genome Browser and incorporated as the standard human NUMT track for the hg38 assembly. Notably, the reverse test was omitted during NUMT identification in this public dataset in order to present a broader set of candidate NUMTs, albeit with a relatively lower confidence level. In total, the track reports 1,072 human NUMTs. To enable visualization, the dataset was converted to bigBed format and implemented as the NuMTs track using a series of UCSC utilities (Figure 4).

**Figure 4:**
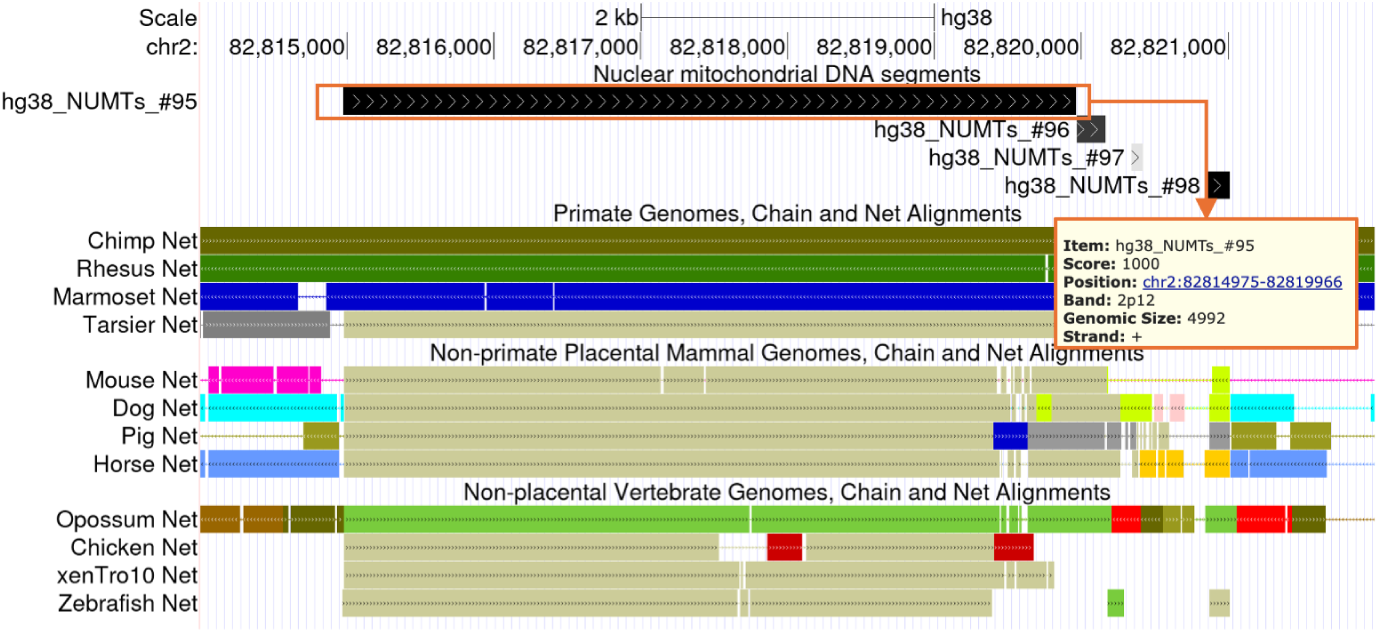
NUMTs UCSC Genome Browser tracks: hg38 reference track generated by our pipeline.Shown is a representative NUMT (hg38 NUMTs #95) on chr2, visualized in the custom NuMTs track. Each NUMT is clickable, revealing details such as genomic position, strand, size, and score. Entries are shaded in varying grayscale to reflect confidence levels, with darker shades indicating higher-confidence NUMTs. Nearby entries (#96–#98) illustrate this gradation. Inter-species alignments are displayed below; beige-colored regions denote segments aligned to the mitochondrial chromosome in the corresponding species.

This NuMTs track provides essential information for each NUMT in the human reference genome, designated as *hg38 NUMTs #*, including genomic co-ordinates, strand orientation, and confidence scores. The arrow direction of each item corresponds to the strand on which the NUMT is aligned. The confidence score for each alignment is calculated as:

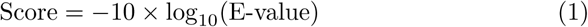

These scores reflect alignment confidence and are visually represented using varying shades of gray, where darker shades indicate stronger confidence. Alignments with score *≥* 100 (i.e. E-value *≤* 1 *×* 10*^−^*^10^) are displayed in black. When a NUMT results from the merging of multiple alignments, the score is taken as the highest among all contributing alignments. While our dataset provides broader genome-wide coverage, the confidence level of individual NUMTs should be interpreted in the context of specific research objectives. We advise users to exercise appropriate caution when referencing individual entries, particularly in downstream analyses that depend on high-confidence annotations.

### 3.4 Survey of NUMTs within mammalian genomes

The optimized NUMT-search pipeline was applied to identify NUMTs in 15 additional mammalian genomes. Summary statistics—including total NUMT counts, total NUMT length, the number of assembled NUMTs blocks and the coverage ratio of NUMTs in the nuclear genome—are presented in Figure 5a. Among the 14 species previously analyzed in other studies, most exhibited higher numbers or longer total lengths of NUMTs using our method, even when restricting the comparison to NUMTs located on assembled chromosomes (Hazkani-Covo, 2022; Uvizl et al., 2024). These results highlight the sensitivity and effectiveness of our approach. An exception was observed in Opossum, where we identified fewer NUMTs than previously reported. This discrepancy is likely due to the absence of scaffold sequences in the opossum genome assembly used in our analysis, which may have excluded some valid NUMT insertions.

**Figure 5:**
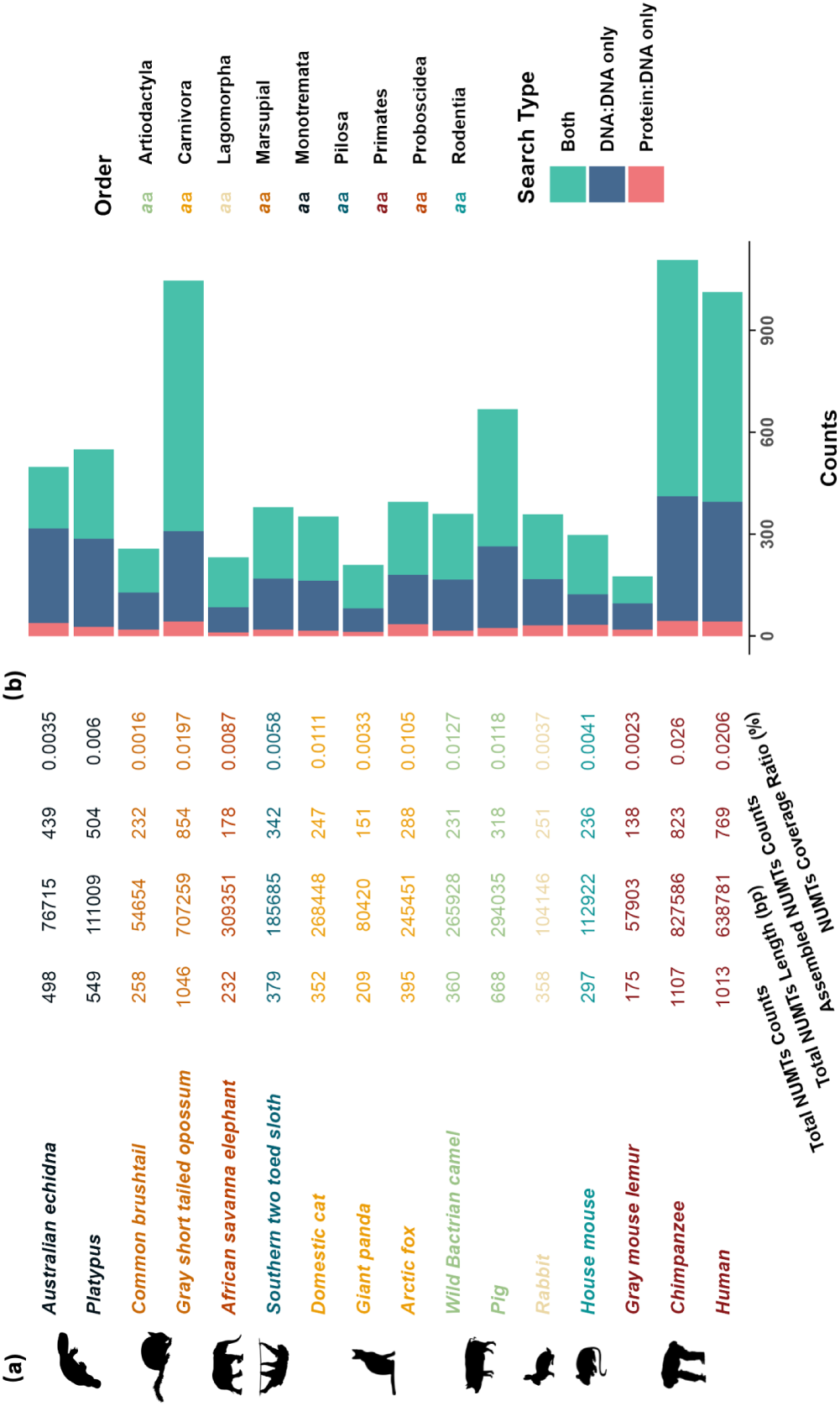
Statistics of NUMTs in 16 mammalian genomes. (a) Summary statistics for each species, including total NUMT counts, total NUMT length (bp), number of assembled NUMTs, and the ratio of total NUMT length to nuclear genome size. Species are grouped and color-coded by mammalian order. (b) NUMTs count distribution by search method. Each stacked bar indicates the number of NUMTs identified by DNA:DNA alignment only (red), protein:DNA alignment only (teal), or both methods (blue). This comparison highlights the complementary contributions of nucleotide- and protein-level searches in different mammalian clades.

To further examine detection patterns, we analyzed the contribution of different alignment methods to NUMT identification using a stacked bar chart (Figure 5b). NUMTs detected by both DNA-to-DNA and DNA-to-protein comparisons comprised the largest fraction, accounting for approximately 40% to 70% of total counts across species. These merged hits, along with NUMTs identified exclusively through DNA-to-DNA alignments, represented the majority of detected NUMTs. Although NUMTs detected solely via DNA-to-protein comparisons were relatively few, typically comprising 5% to 10% of the total, they highlight the importance of incorporating protein-level alignments to improve detection sensitivity, particularly for highly diverged insertions.

We also analyzed the length distribution of NUMTs across species, as shown in Figure 6, which includes the median NUMT length for each genome. Most NUMTs were shorter than 1 kb, comprising 76% to 99% of total NUMT counts per species. Notably, the longest NUMT identified was found in the cat genome, spanning 64,876 bp. This element is consistent with a previously reported large-scale mitochondrial insertion in the domestic cat, comprising a tandem array of a 7.9-kb mitochondrial fragment located on chromosome D2 (Lopez et al., 1994).

**Figure 6:**
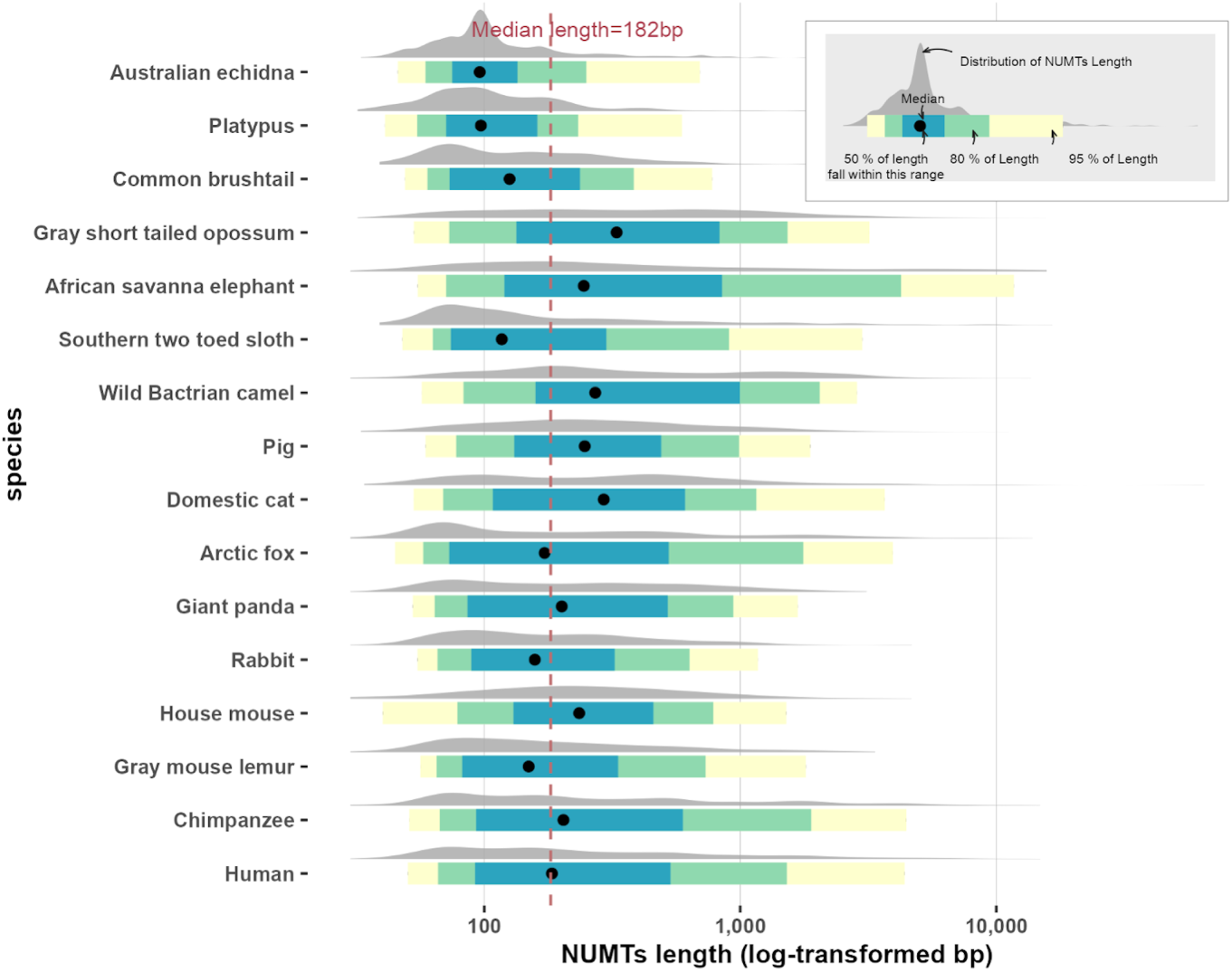
Distribution of NUMT lengths across 16 mammalian species. Ridgeline plots display the log-transformed length distributions of NUMTs in each species. Colored intervals represent the 50%, 80%, and 95% quantile ranges, with black dots indicating medians. The vertical dashed line marks the overall median NUMT length.

### 3.5 Ancient NUMT insertions across mammalian clades

By comparing ancient NUMTs and orthologous segments in 120 pair-wise alignments, we established the NUMTs orthologous network among 16 species (see Supplementary Table S2). The insertion time of these ancient NUMTs was inferred based on their presence at nodes within the phylogenetic tree of selected mammalian genomes, as established by Timetree.org. Since we mainly focus on ancient NUMTs, the NUMTs orthology network is based on NUMTs inserted before the last common ancestor of genomes in the same mammalian order.

Most of the ancient NUMTs in mammals were inserted within the last 100 million years, with only a few appearing to have been integrated near the root of the mammalian phylogenetic tree. A 63-bp camel NUMT with an E-value of 0.039, identified via the DNA-to-protein comparison, overlaps with orthologous regions in nine other mammalian genomes—including echidna. However, it does not overlap with any annotated NUMTs in these genomes. Ancient NUMTs, by nature, are more challenging to detect compared to recent insertions due to the accumulation of mutations and genomic rearrangements over time. The lower confidence reflected in the E-value shows an inevitable limitation when searching for highly degraded NUMTs that have persisted for such a long evolutionary time.

A total of 189 ancient orthologous NUMTs were identified in the human genome, among which 90 were estimated to have been inserted at least 100 million years ago (Figure 7). The lowest E-value among them was 6.3 *×* 10*^−^*^100^, for a NUMT that overlaps with three other genomes, including that of the sloth. To our knowledge, no previous studies have reported ancient NUMTs predating the common ancestor of Boreoeutheria (Uvizl et al., 2024). The identification of such deeply conserved elements substantially extends the detectable timeline of NUMT insertion events.

**Figure 7:**
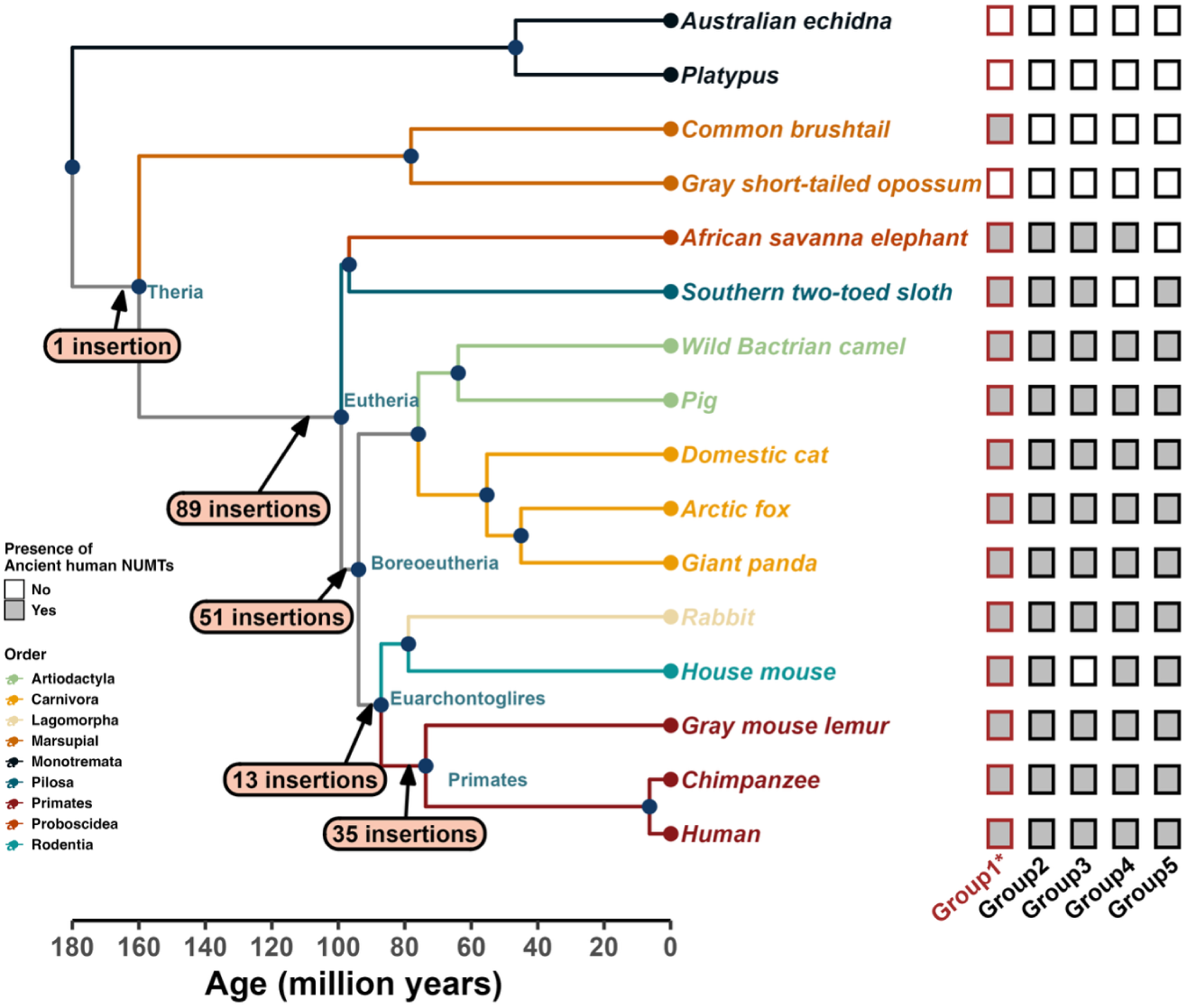
The most ancient NUMTs in the human genome. The left panel shows the phylogenetic tree of 16 mammalian species used to trace ancient human NUMTs, with major divergence nodes annotated with the number of inferred NUMT insertions. Branches and species names are color-coded by mammalian order, as indicated in the legend below. The right panel summarizes the presence or absence of the five most ancient NUMT groups (Group 1–5) in the human genome. Filled black squares indicate orthologous NUMTs present in both human and the corresponding species. Red-bordered squares denote putative NUMTs—highly degenerated NUMTs not detectable directly in the human genome but inferred through conserved alignments with other species.

We selected the 11 most conserved NUMTs, shared by humans and more than 10 other species, and grouped them into five categories based on their presence in inter-genome alignments (Table 1). Notably, the most ancient NUMT (Group 1) was inferred to have originated at least 160 million years ago. This is a therian NUMT inferred from a conserved NUMT in the brushtail possum with an E-value of 0.00092, which overlaps with orthologous regions in 11 mammalian genomes. This human NUMT was not directly detected by our NUMT detection pipeline, likely due to extreme sequence degeneration. However, by leveraging the alignment with the possum genome, we were able to infer its presence as a putative ancient human NUMT. To further confirm the conservation of this brushtail NUMT, we conducted additional searches for NUMTs in two other marsupials, wallaby and Tasmanian devil, as well as orthologous NUMTs searches between brushtail and these species. Results show that this brushtail NUMT overlaps with a NUMT in the wallaby genome (E-value = 0.018) and another in the Tasmanian devil genome (E-value = 2.6 *×* 10*^−^*^26^), supporting with high confidence that the insertion likely occurred in the common ancestor of therian mammals. This NUMT originates from a mitochondrial tRNA region, consistent with prior findings that tRNA-derived NUMTs exhibit slower evolutionary rates than adjacent numtDNA sequences (Hoser et al., 2020). These results raise intriguing possibilities. Ancient tRNA-derived NUMTs may serve as sources of novel regulatory elements or influence nuclear genome stability through long-term conservation and integration. Further investigation is warranted to explore the potential functional consequences of such deeply conserved NUMTs.

**Table 1:** NUMTs data description across chromosomes.

| Chromosome | ChromStart | ChromEnd | Description | Group |
| --- | --- | --- | --- | --- |
| *chr2* | 118055298 | 118055357 | potential | 1 |
| *chr2* | 40955198 | 40955359 | human NUMTs assembled#63 | 2 |
| *chr3* | 71949580 | 71949689 | human NUMTs assembled#146 | 2 |
| *chr7* | 116306306 | 116306513 | human NUMTs assembled#343 | 2 |
| *chr16* | 17992090 | 17992363 | human NUMTs assembled#608 | 2 |
| *chr4* | 123837642 | 123837724 | human NUMTs assembled#222 | 3 |
| *chr5* | 37164410 | 37164493 | human NUMTs assembled#244 | 3 |
| *chr7* | 37518741 | 37518852 | human NUMTs assembled#324 | 3 |
| *chr11* | 37541525 | 37543000 | human NUMTs assembled#501 | 3 |
| *chr9* | 7511591 | 7511719 | human NUMTs assembled#414 | 4 |
| *chr5* | 160339468 | 160339652 | human NUMTs assembled#278 | 5 |

Although functional characterization lies beyond the scope of this study, we consider the ancient NUMTs identified here to be a valuable foundation for future investigations into organellar DNA-derived elements within the nuclear genome. Among these, short ancient NUMTs may warrant particular attention. Short NUMT insertions represent the majority of nuclear organellar DNA fragments in eukaryotic genomes (Richly and Leister, 2004) and have been reported as promising candidates for recruitment into nuclear reading frames, with the potential to evolve into novel exons (Noutsos et al., 2007).

While not all ancient NUMTs are functionally active, their evolutionary significance should not be underestimated. NUMTs shared across multiple species provide evidence of insertion events predating lineage divergence, thereby serving as molecular markers for reconstructing deep phylogenetic relationships. Furthermore, the persistence and genomic distribution patterns of ancient NUMTs offer insights into the historical dynamics of mitochondrial–nuclear interactions (Bensasson et al., 2001). These sequences serve as genomic records of the long-term co-evolution between mitochondrial and nuclear genomes over tens of millions of years.

## 4 Conclusions and Prospects

Detecting highly degraded NUMTs remains challenging due to the limitations of conventional search strategies, which hinder our understanding of their potential influence on the genome and organism. This study proposes a novel detection pipeline using a refined sequence comparison method to overcome this limitation. The pipeline is designed to comprehensively identify all NUMTs within the nuclear genome, with a particular emphasis on recovering ancient NUMT alignments that might otherwise be obscured by background noise. This exhaustive approach comes at the cost of significantly increased computational time compared to conventional methods that rely on local alignment tools.

Using this pipeline, we completed the annotation of NUMTs in the human genome assembly GRCh38, which has now been implemented in the UCSC Genome Browser. Analysis of NUMT orthology across 16 mammalian genomes revealed many ancient NUMTs that have not been previously reported. While the majority of ancient NUMTs identified in this study appear to have been inserted into the nuclear genome of the common ancestor of Eutherian mammals, even older NUMTs may be discovered with expanded analyses across genomes from more diverse orders.

When investigating a few extremely ancient orthologous NUMTs, a major challenge was distinguishing genuine conservation across species from cases where a NUMT happens to align by chance to a highly conserved region in a single genome. Factors such as E-value, insertion sites in the host genome, and the presence of a NUMT exclusively in one species should be carefully considered. One concern about the findings was that even with strict confidence thresholds and measures to minimize random alignments, false positives in NUMT detection cannot be entirely ruled out. Although it may be impossible to conclusively prove that these NUMTs are truly ancient, broader comparative analysis across more species could provide stronger support for their orthology.

In summary, this study expands our understanding of NUMTs and illustrates how tailored computational strategies can help detect ancient and subtle genomic signals. These findings are expected to inspire further research into the evolutionary and functional significance of NUMTs, as well as other relics of ancient genomic processes.

## Acknowledgments

We thank members of the Frith and Asai laboratories and Silvia Rodriguez for helpful discussions, feedback, and support throughout the course of this work. This work was supported by JST SPRING, Grant Number JPMJSP2108 and by the Japan Science and Technology Agency, Grant Number JPMJCR21N6.

